# Decreasing ZAP Poly(ADP-ribose) Binding Enhances Antiviral Activity Against Alphaviruses

**DOI:** 10.1101/2024.08.30.610567

**Authors:** Shang-Jung Cheng, Diane E. Griffin, Anthony K. L. Leung, Rachy Abraham

## Abstract

Zinc finger antiviral protein (ZAP), also known as PARP13, is an antiviral factor effective against several virus families. ZAP contains three major protein domains: N-terminal zinc fingers, central WWE domains, and a C-terminal inactive ADP-ribosyltransferase domain, which is present only in the longer isoform ZAPL. While both the zinc finger and ADP-ribosyltransferase domains inhibit viral replication, the role of the WWE domains remains unclear. WWE domains bind poly(ADP-ribose) (PAR). In this study, we focus on ZAPS, the shorter isoform comprised of the minimal unit that confers antiviral activity and lacking the ADP-ribosyltransferase domain, to investigate how PAR-binding contributes to antiviral defense against alphaviruses. We identified the Y659 residue on the second WWE domain as essential for PAR-binding both in vitro and in cells. When infected with Chikungunya or Sindbis virus, cells with the Y659A mutant of ZAPS had reduced formation of replication complexes, decreased levels of viral genomic and subgenomic transcripts, and fewer infectious virions released compared to cells with the unmutated ZAPS, suggesting enhanced antiviral activity of the PAR-binding mutant. These findings suggest that inhibiting PAR-binding in ZAPS could potentiate host antiviral functions, offering a novel therapeutic strategy against alphavirus infections.

## IMPORTANCE

Alphaviruses are mosquito-borne RNA viruses that cause rash, arthritis, and neurological disease. Despite the ongoing threat of outbreaks, there are no licensed treatments available for any alphavirus. Effective control of these virus infections hinges on a deeper understanding of the molecular targets—both viral and host factors—that regulate viral replication. In this study, we uncover a novel mechanism to enhance the antiviral function of a key host factor by inhibiting its ability to bind poly(ADP-ribose) (PAR)—often referred to as the “third nucleic acid” because of its similarity to DNA and RNA and its critical role in many fundamental processes.

Our findings show that PAR-binding activity is crucial for viral replication and the production of infectious virus particles. This mechanism is conserved across both Chikungunya and Sindbis viruses, highlighting a novel target for developing anti-alphaviral therapies by inhibiting the PAR-binding function of ZAPS.

## OBSERVATION

### ZAPS Lacking PAR Binding Exhibits Enhanced Antiviral Activity Against Alphavirus

Zinc-finger antiviral protein, present in two major isoforms (ZAPL and ZAPS), plays a critical role in host defense against viral infections, demonstrating broad antiviral activity across multiple virus families (1), including alphaviruses with pandemic potential but limited drug targeting options. ZAP, also known as PARP13, is one of the 17 human members of the ADP-ribosyltransferase family responsible for ADP-ribosylation, including generating the polymeric form poly(ADP-ribose) or PAR. The shorter ZAPS isoform, generated by alternative splicing, shares the same N-terminal zinc fingers and central tandem PAR-binding WWE domains as the longer ZAPL isoform (2, 3), but ZAPL additionally contains a catalytically inactive ADP-ribosyltransferase domain at its C-terminus **(Fig. 1A)**. The zinc fingers in ZAP recognize CpG-rich viral RNA and recruit RNA degradation machinery to inhibit viral replication (4). These zinc fingers also interact with the cofactor protein TRIM25 to boost antiviral activity (5), while the ADP-ribosyltransferase domain engages with viral proteins to suppress replication via proteasome degradation (6). However, the antiviral function of the WWE domains remains largely unclear. A recent study reported that mutating Q668 in the WWE domain leads to a loss of PAR-binding and weakens the antiviral ability of ZAPS against an engineered HIV variant with high CG content (3). However, it is unclear whether such a phenomenon extends to other viruses in their native contexts. Given ZAPS’s simpler domain structure and demonstrated sufficiency for antiviral activity, we aimed to use it to decipher the role of PAR-binding in WWE domain activity against alphaviruses.

**Figure 1.**
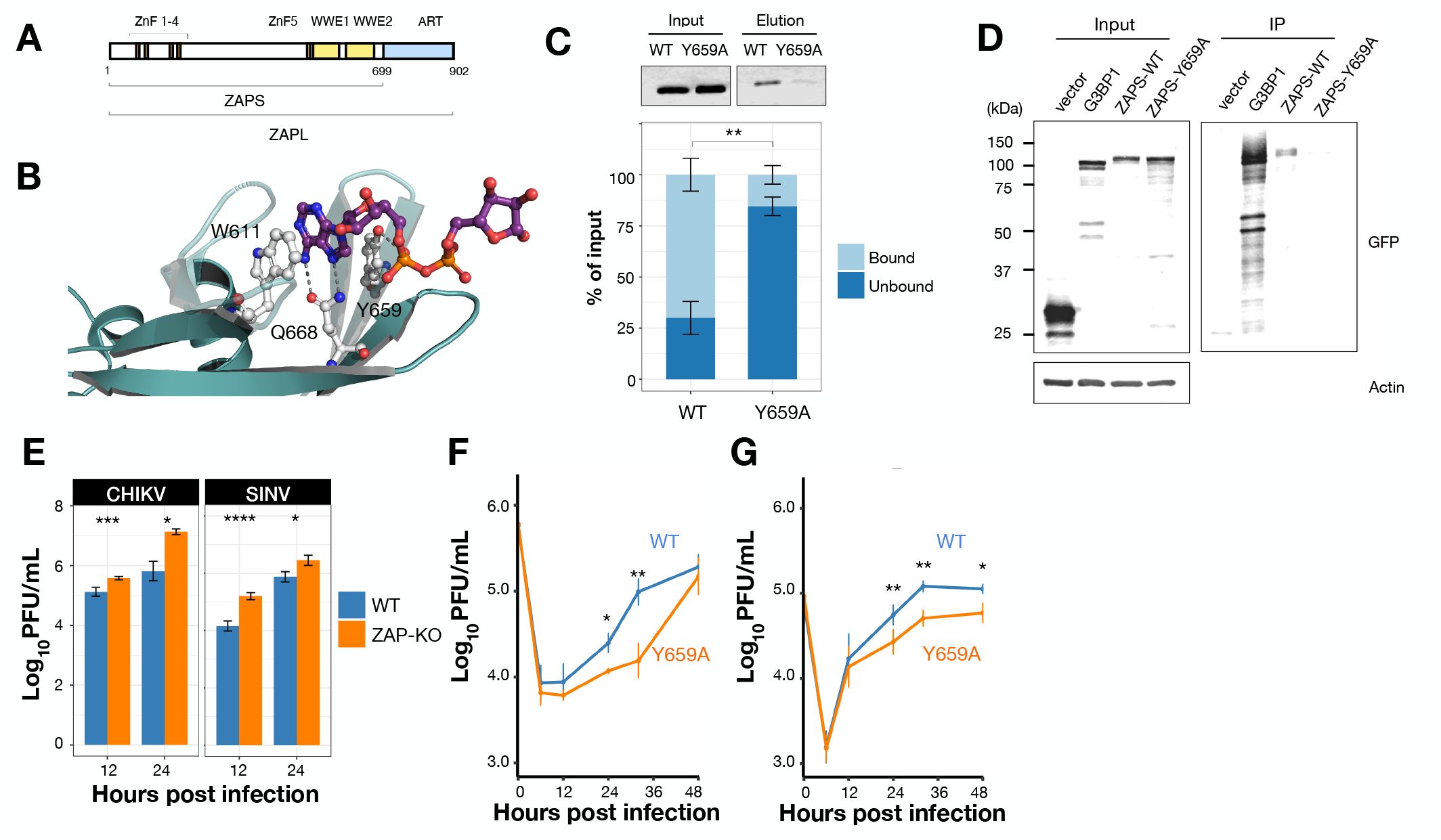
| ZAPS Lacking PAR Binding Exhibits Enhanced Antiviral Activity Against Alphavirus. A. Domain structures of the two major ZAP isoforms. B. Similar to the established PAR-binding residues W611 and Q688, Y659 is located in the ADP-ribose-binding pocket of ZAPS (PDB: 7tgq). Dash lines indicate potential hydrogen bonds with ADP-ribose. C. Purified wild-type or Y659A mutant ZnF-WWE1-WWE2 fragments were incubated with biotinylated 11-mer PAR and pulled down with streptavidin beads. Proteins eluted from the beads were regarded as “bound” fractions and, together with the unbound fraction, were analyzed by SDS-PAGE. The percentage of eluted proteins was normalized to input and mean ± standard error was graphed by R studio (ggplot2). D. Cell lysate expressing GFP-tagged vector (negative control), G3BP1(positive control), wild-type ZAPS, or Y659A was incubated with biotinylated 11-mer PAR, followed by streptavidin beads pull down and analyzed by SDS-PAGE. E. HeLa wild-type and ZAP knockout cells were infected with CHIKV 181/25 or SINV TE strains at an MOI 1. Supernatant was collected after 12 and 24 hpi for plaque assay to determine viral titer. Mean ± standard deviation was graphed by R studio (ggplot2). F. HeLa ZAP knockout cells were transfected with GFP-tagged wild-type or Y659A ZAPS for 24 h. Cells were infected with CHIKV 181/25 (MOI = 1). Supernatants were collected 6, 12, 24, 32, and 48 hpi for plaque assay to determine viral titers. Mean ± standard deviation was graphed by R studio (ggplot2). G. HeLa ZAP knockout cells were transfected with GFP-tagged wild-type or Y659A ZAPS for 24 h. Cells were infected with SINV TE (MOI = 1). Supernatants were collected 6, 12, 24, 32, and 48 hpi for plaque assay to determine viral titers. Mean ± standard deviation was graphed by R studio (ggplot2). All results were obtained from three independent experiments. Statistical analysis was determined by a two-tailed paired Student t-test. *P < 0.05, ** P < 0.01, ***P < 0.001, ****P < 0.0001.

Biochemical and structural studies suggest that the second of the tandem WWE domains plays a significant role in PAR-binding (2, 3). Sequence alignment with the well-characterized WWE domain of ubiquitin E3 ligase RNF146 reveals that the Y659 residue in PARP13 is conserved with the key PAR-binding site Y144 of RNF146 (3). Structurally, Y659 is located in the ADP-ribose binding cavity, similar to the established PAR-binding residues, W611 and Q668, in ZAPS (2, 3) (**Fig. 1B**), suggesting its potential role in binding to PAR. To test this hypothesis, we incubated recombinant wild-type and Y659A protein fragments with 11-mer biotinylated PAR, followed by pulldown with streptavidin beads. After elution and SDS-PAGE analysis, we found that wild-type protein bound to PAR three times more than the Y659A mutant in vitro **(Fig. 1C)**. In cell lysates expressing GFP-tagged full-length ZAPS and the Y659A mutant, Y659A completely lost its binding ability when incubated with biotinylated PAR **(Fig. 1D)**. Taken together, these data indicate that Y659 is essential for ZAP to bind PAR in vitro and in cells.

To assess whether the PAR-binding is critical for the antiviral activity of ZAPS in infected cells, we first validated its activities against alphaviruses by comparing CHIKV and SINV replication at an MOI of 1 in HeLa ZAP knockout and parental wild-type (WT) cells. Virus production quantified by plaque assay in Vero cells indicated that replication of both alphaviruses was restricted in wild-type HeLa cells compared to the ZAP knockout counterparts **(Fig. 1E)**. Next, we expressed GFP-tagged ZAPS wild-type and the PAR binding mutant Y659A in ZAP knockout HeLa cells for 24 h. CHIKV titers from Y659A-expressing cells were significantly lower than those in wild-type cells, particularly at the later time points, 24 h and 32 h post infection (hpi) **(Fig. 1F)**. A similar pattern was observed for SINV **(Fig. 1G)**, suggesting that the enhanced antiviral activity of the mutant with decreased PAR-binding is conserved between alphaviruses.

### PAR-Binding Ability of ZAPS Is Critical for Replication Complex Formation and Viral Transcript Synthesis

To further explore how PAR-binding ability by ZAPS affects the viral replication cycle, we compared the replication complexes formed in cells expressing wild-type or PAR-binding mutant Y659A by staining for dsRNA, which is indicative of replication complexes, using flow cytometry. Compared to the negative control cells expressing the GFP-tagged AID (auxin-inducible degron), cells expressing either the wild-type or Y659A constructs had less dsRNA staining **(Fig. 2A)**. Consistently, the dsRNA level in infected Y659A cells was even lower than in wild-type-expressing cells at 24 hpi **(Fig. 2A)**, suggesting that cells expressing the PAR-binding ZAPS mutant are less supportive of viral replication complex formation than cells with WT ZAPS.

**Figure 2.**
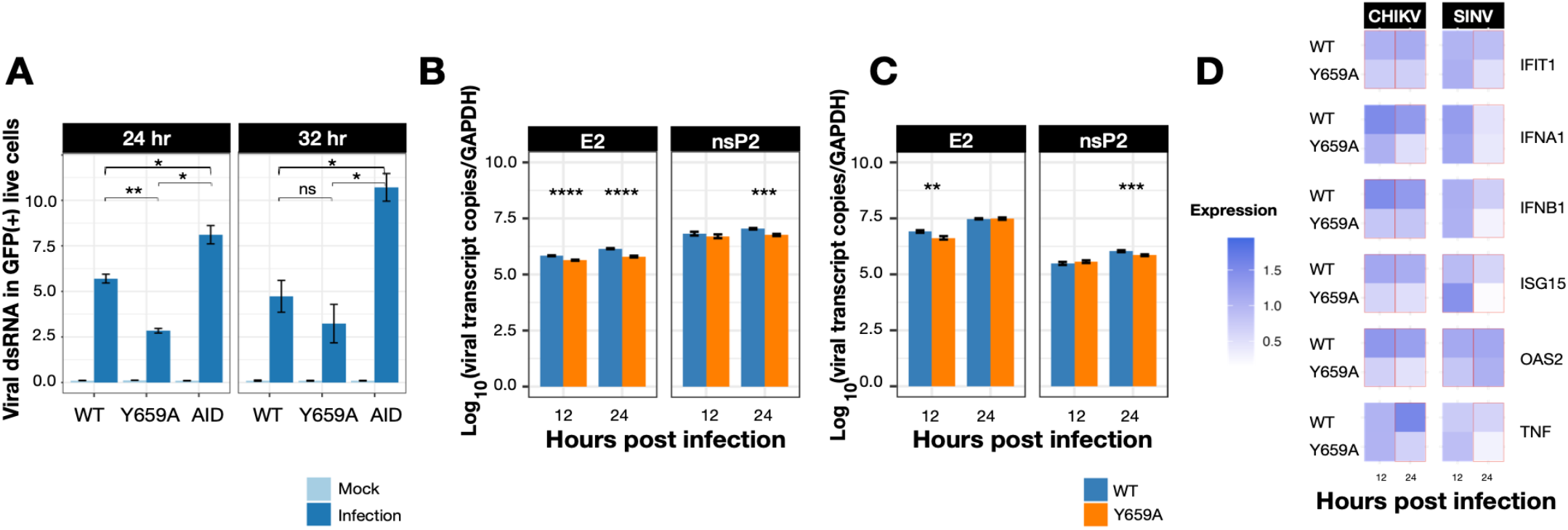
| PAR-Binding Ability of ZAPS Is Critical for Replication Complex Formation and Viral Transcript Synthesis. A. GFP-tagged wild-type full-length ZAPS, Y659A, and AID were transfected into HeLa ZAP knockout cells and infected with CHIKV 181/25 (MOI = 10). Cells were harvested after 24 or 32 h for dsRNA staining (mouse anti-J2, Scicons) and analyzed by flow cytometry. The percentage of viral dsRNA in GFP-positive cells was measured and graphed using R Studio (ggplot2). B. HeLa ZAP knockout cells were transfected with GFP-tagged wild-type or Y659A ZAPS for 24 h. Cells were infected with CHIKV 181/25 (MOI = 1). RNA was isolated from infected cells and reverse-transcribed to cDNA. qRT-PCR was performed to quantify viral transcripts, E2 and nsP2. Values were back-calculated to copy numbers and normalized to GAPDH levels. Mean ± standard error was graphed by R studio (ggplot2). C. HeLa ZAP knockout cells were transfected with GFP-tagged wild-type or Y659A ZAPS for 24 h. Cells were infected with (C) SINV TE (MOI = 1). RNA was isolated from infected cells and reverse-transcribed to cDNA. qRT-PCR was performed to quantify viral transcripts, E2 and nsP2. Values were back-calculated to copy numbers and normalized to GAPDH levels. Mean ± standard error was graphed by R studio (ggplot2). D. RNA was isolated from infected and mock-infected cells and reverse-transcribed to cDNA. qRT-PCR was performed to quantify the expression of host interferon-stimulated genes. Values of each gene were normalized to ACTB level and mock condition. A tile graph was used to visualize the mean values of gene expression level (value of 2 ^-ΔCT^). Genes with statistical differences of p < 0.05 between the wild-type and Y659A mutant were framed by red lines. All results were obtained from three independent experiments. Statistical analysis was determined by a two-tailed paired Student t-test. *P < 0.05, ** P < 0.01, ***P < 0.001, ****P < 0.0001, ns: not significant.

To determine whether differences in replication complex abundance affect the synthesis of viral RNAs, we measured genomic and subgenomic RNA at early and late stages of alphavirus infection by qRT-PCR. The transcripts for both structural protein E2 (subgenomic RNA) and nonstructural protein nsP2 (genomic RNA) were slightly higher in cells expressing wild-type ZAPS compared to cells expressing the Y659A mutant at either the early or late stages of alphavirus infection **(Fig. 2B-C)**. These data suggest that the Y659A mutant impairs the synthesis of viral transcripts, thereby restricting alphavirus replication.

Because ZAPS also mediates host innate immune responses by modulating RIG-I activation and enhancing interferon signaling during viral infection (7, 8), the PAR-binding mutant might further increase interferon pathway responses, leading to greater virus restriction. To test this hypothesis, we performed qRT-PCR to quantify the expression of interferons (IFNA1 and IFNB1) and a panel of interferon-stimulated genes (ISGs) during CHIKV and SINV infection. Although these genes were induced, the level was low, likely due to ZAPS-mediated suppression (7).

Contrary to our hypothesis, both interferons, ISGs, including IFIT1, ISG15, OAS2, and pro-inflammatory TNF, were more strongly induced in wild-type ZAPS-expressing cells perhaps due to higher virus replication in wild-type cells **(Fig. 2D)**.

## DISCUSSION

Previous alphavirus studies of ZAP function showed that the N-terminal zinc fingers target specific CpG-rich regions on viral nsP2 (9) or alter cellular localization (10) to inhibit viral replication or translation. In this study, we identified the independent role of PAR-binding in the WWE domain in regulating viral replication. The Y659A mutation that decreased PAR binding enhanced ZAPS antiviral activity. The increased protection of the PAR-binding mutant against alphavirus infection, though opposite to its effect in the engineered high-CG-content HIV variant, suggests that the role of PAR binding may vary depending on the virus and its replication mechanisms. Because we observed differences in alphavirus infection between cells expressing wild-type ZAPS and Y659A mutant ZAPS in replication complex formation and viral transcript synthesis, it is likely that PAR binding affects virus infection at or prior to the initiation of replication.

Alphaviruses have an intriguing relationship with ADP-ribosylation. Alphaviruses encode highly conserved macrodomains in the nonstructural protein nsP3, which bind to and hydrolyze ADP-ribose on modified proteins. Both the binding and hydrolase activities are essential for viral replication and virulence in mice (11). Recently, our group identified that this macrodomain is also crucial for the disassembly of stress granules—cytoplasmic condensates formed during viral infection that are critical for antiviral function and often targeted by viruses for disassembly through various mechanisms. For example, the hydrolase activity of the CHIKV macrodomain specifically removes ADP-ribosylation from stress granule proteins, including the core component G3BP1, and disassembles virus-induced stress granules (12). Stress granules are notably enriched with PAR, where ZAPS is also located during viral infection (10). Therefore, if the PAR-binding mutant of ZAPS delays stress granule disassembly, it may sequester essential factors for viral replication and translation, slowing virus production. Our findings thus suggest that inhibiting the PAR-binding of ZAPS could enhance host antiviral function, guiding future therapeutic strategies against alphavirus infections.

## Material and Methods

### Cell Culture and plasmids

The HeLa ZAP knockout cell line and eGFP-tagged wild-type ZAPS construct were gifts from Dr. Paul Chang. eGFP-tagged AID construct was shared by Dr. Andrew Holland (Johns Hopkins University). Cell lines were maintained in DMEM supplemented with 10% heat-inactivated FBS (Gibco, Life Technologies) at 37 °C in a 5% CO_2_ incubator. ZAPS Y659A mutant construct was prepared by site-directed mutagenesis and confirmed by sequencing.

### Viruses

CHIKV vaccine strain 181/25 virus was rescued in BHK21 cells from transfected RNA, in vitro transcribed from a full-length cDNA clone (gift from Naomi Forrester, University of Texas Medical Branch, Galveston, TX). Viral stocks were grown in BHK21 cells and titrated in Vero cells. SINV TE strain virus was rescued in BHK21 cells from transfected RNA, invitro transcribed from a full-length cDNA clone. Viral stocks and titration were performed in BHK21 cells.

### ZAP Overexpression and virus infection

HeLa ZAP knockout cells were seeded and transfected either with eGFP-tagged wild-type ZAPS (WT) or eGFP-tagged ZAPS Y659A (Y659A) or eGFP-tagged AID (AID) construct using Lipofectamine 2000 (Invitrogen). At 24 h post-transfection, the cells were mock infected or infected with CHIKV or SINV at MOIs of 1 or 10 and incubated for an hour at 37 °C. The media was replaced with DMEM supplemented with 1% heat-inactivated FBS and incubated for different hours post infection at 37 °C in a 5% CO_2_ incubator.

Supernatant collected at different hours post infection (hpi) was serially diluted and added to Vero monolayers and incubated for 1 h. The inoculum was removed and overlaid with a media having equal proportion of 2X Modified Eagle Medium (Gibco) supplemented with 1% heat-inactivated FBS media and 2% Carboxymethyl cellulose (Sigma) and incubated at 37 °C for 48 h. Cells were fixed with 10% formaldehyde in PBS and stained with 0.02% crystal violet for plaque counting. Data was quantified as PFU per milliliter and visualized in line graphs.

### Quantification of dsRNA using flow cytometry

HeLa ZAP knockout cells were seeded and transfected with Lipofectamine 2000, followed by mock infection or infection at an MOI of 10 with CHIKV. At 24 and 32 hpi, cells were collected and stained with live/dead staining (Invitrogen) on ice for 30 min. Cells were fixed with 2% paraformaldehyde and were permeabilized with 0.2% Triton in FACS buffer (PBS with 0.4% 500 mM EDTA and 0.5% BSA). dsRNA was stained with J2 mouse monoclonal antibody (1:1,000; Scicons) for 1 h on ice, followed by PE-conjugated goat anti-mouse IgG secondary antibody staining. Samples were analyzed on a FACSCanto flow cytometer. Bar graph was used to visualize the averaged percentage of dsRNA in GFP-positive live cells.

### Quantification of Viral RNA transcripts

Total RNA of mock-infected and infected cells was isolated with the High Pure RNA Tissue kit (Roche) and 500 ng of RNA was reverse transcribed to cDNA using High-Capacity cDNA Synthesis Kit (Applied Biosystems). qRT-PCR was performed with Taqman probe specific for viral nsP2 and E2 region or primers against these genes using Universal PCR Master Mix (Applied Biosystems) (see Table 1: Details of primer and probe sequences). Plasmid constructs containing CHIKV E2/nsP2, SINV E2/nsP2, or Rodent GAPDH were 10-fold serial-diluted for standard curves of each transcript, which were amplified simultaneously with sample cDNAs. Ct values of transcripts were converted to copy numbers based on corresponding standard curves, followed by GAPDH normalization. Data was visualized as the log value of the average viral transcript copies per 10^6^ copies of GAPDH. All reactions were run on the Applied Biosystems 7500 Real-time PCR machine under the following conditions: 50 °C for 2 min, 95 °C for 10 min, 95 °C for 15 s, and 60 °C for 1 min for 40 cycles and were analyzed with 7500 Software version 2.0.5 (Applied Biosystems).

**Table 1.**
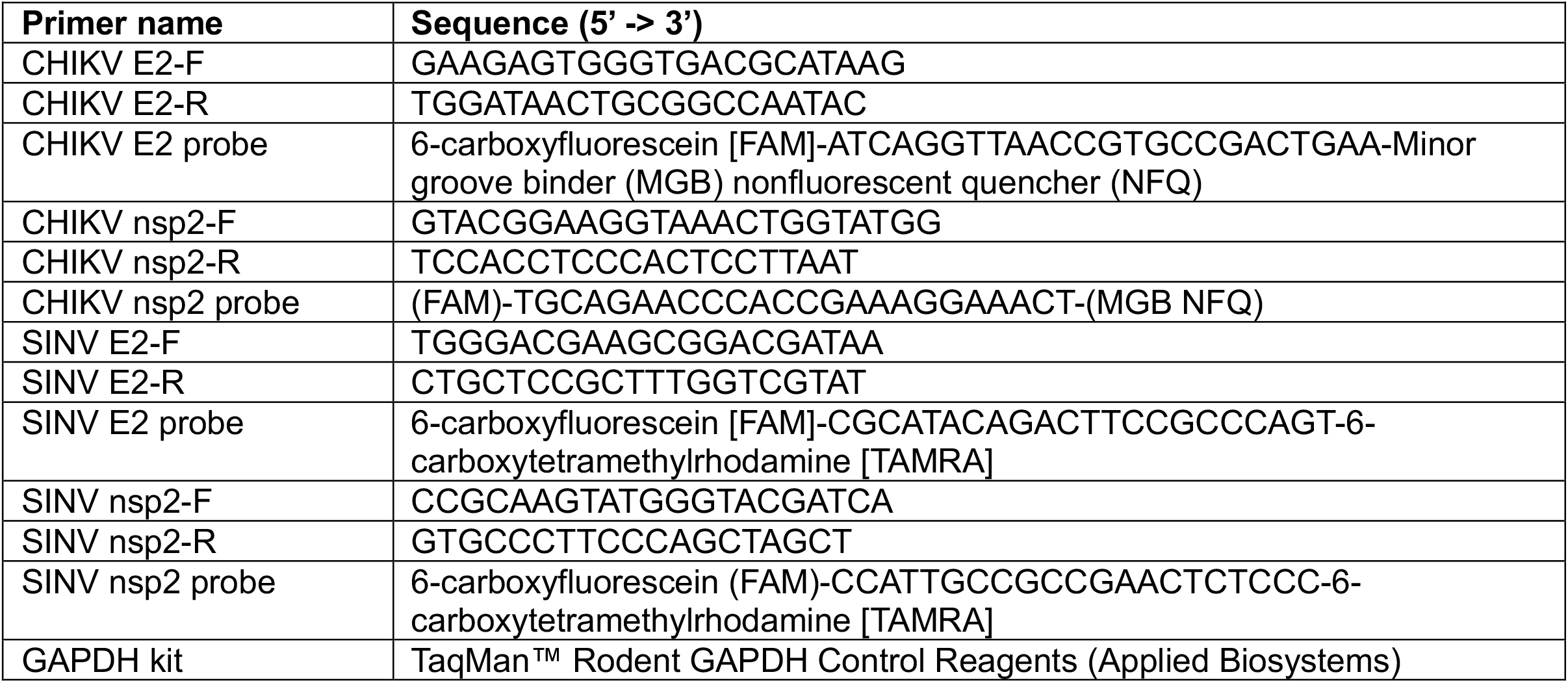
Taqman primers/probes.

### qRT-PCR for Host Gene Expression

Total RNA of mock-infected and infected cells was isolated, and reverse transcribed to cDNA, as previously described. qRT-PCR was performed by using PowerUp SYBR Green Master Mix (Applied Biosystems) and quantifying with ΔΔC_T_ method (Table 2: Details of primer sequences). Relative gene expression was normalized by both mock-infected samples at the corresponding time point and *ACTB* level. Values were converted to 2^-ΔΔCT^ to show expression level and were visualized by tile graph. All reactions were run on the Applied Biosystems 7500 Real-time PCR machine under the following conditions: 50 °C for 2 min, 95 °C for 10 min, 95 °C for 15 s, and 60 °C for 1 min for 40 cycles and were analyzed with 7500 Software version 2.0.5 (Applied Biosystems).

**Table 2.**
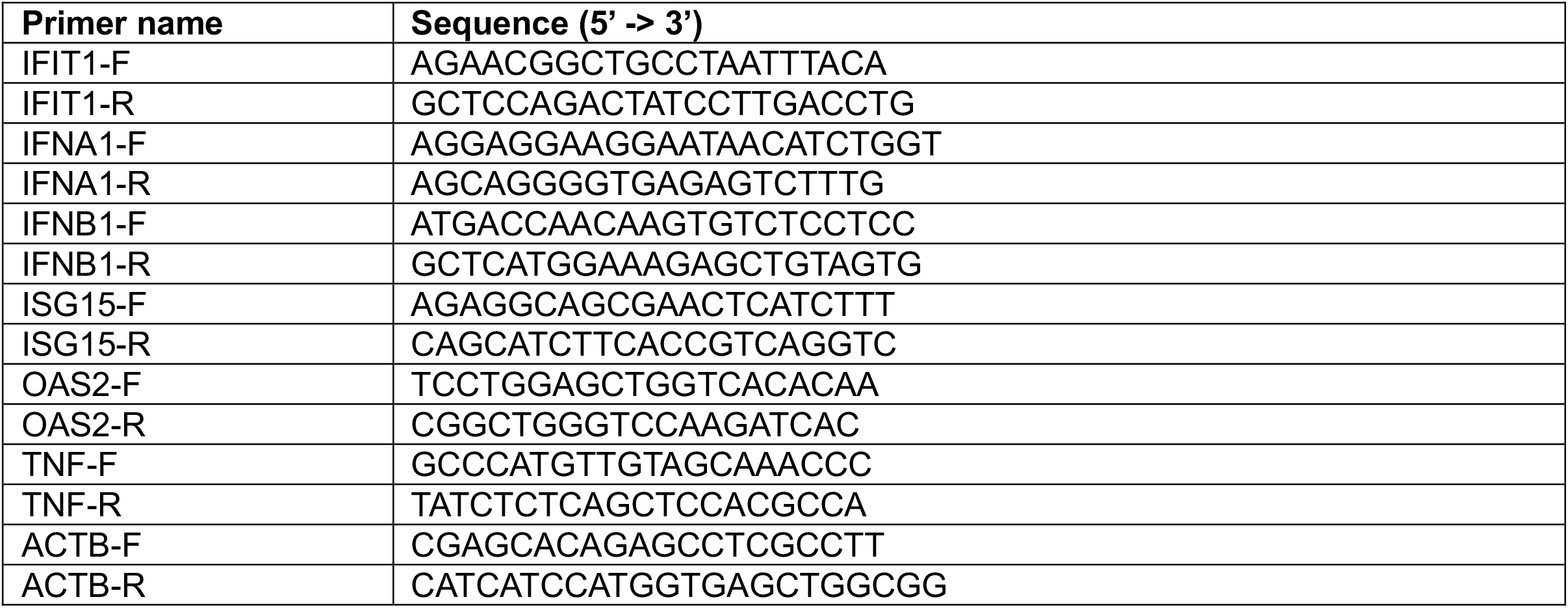
Primer sequence.

### Statistics and data visualization

Differences between groups were determined by a paired, two-tailed Student *t*-test. All statistical analysis was perfomred using ggplot2 (R Studio); results are shown as means ± SD or means ± SE (details in figure legend).

## REFERENCES

1. Ficarelli M, Neil SJD, Swanson CM. 2021. Targeted Restriction of Viral Gene Expression and Replication by the ZAP Antiviral System. Annu Rev Virol 8:265–283. 10.1146/annurev-virology-091919-104213.

2. Kuttiyatveetil JRA, Soufari H, Dasovich M, Uribe IR, Mirhasan M, Cheng S, Leung AKL, Pascal JM. 2022. Crystal structures and functional analysis of the ZnF5-WWE1-WWE2 region of PARP13/ZAP define a distinctive mode of engaging poly(ADP-ribose). Cell Rep 41:111529. S2211-1247(22)01385-7 [pii].

3. Xue G, Braczyk K, Goncalves-Carneiro D, Dawidziak DM, Sanchez K, Ong H, Wan Y, Zadrozny KK, Ganser-Pornillos BK, Bieniasz PD, Pornillos O. 2022. Poly(ADP-ribose) potentiates ZAP antiviral activity. PLoS Pathog 18:e1009202. 10.1371/journal.ppat.1009202[doi].

4. Zhu Y, Chen G, Lv F, Wang X, Ji X, Xu Y, Sun J, Wu L, Zheng Y, Gao G. 2011. Zinc-finger antiviral protein inhibits HIV-1 infection by selectively targeting multiply spliced viral mRNAs for degradation. Proc Natl Acad Sci U S A 108:15834–15839. 10.1073/pnas.1101676108.

5. Li MMH, Lau Z, Cheung P, Aguilar EG, Schneider WM, Bozzacco L, Molina H, Buehler E, Takaoka A, Rice CM, Felsenfeld DP, MacDonald MR. 2017. TRIM25 Enhances the Antiviral Action of Zinc-Finger Antiviral Protein (ZAP). PLoS Pathog 13:e1006145. 10.1371/journal.ppat.1006145.

6. Liu C, Zhou L, Chen G, Krug RM. 2015. Battle between influenza A virus and a newly identified antiviral activity of the PARP-containing ZAPL protein. Proc Natl Acad Sci U S A 112:14048– 14053. 10.1073/pnas.1509745112.

7. Schwerk J, Soveg FW, Ryan AP, Thomas KR, Hatfield LD, Ozarkar S, Forero A, Kell AM, Roby JA, So L, Hyde JL, Gale MJ, Daugherty MD, Savan R. 2019. RNA-binding protein isoforms ZAP-S and ZAP-L have distinct antiviral and immune resolution functions. Nat Immunol 20:1610–1620. 10.1038/s41590-019-0527-6.

8. Hayakawa S, Shiratori S, Yamato H, Kameyama T, Kitatsuji C, Kashigi F, Goto S, Kameoka S, Fujikura D, Yamada T, Mizutani T, Kazumata M, Sato M, Tanaka J, Asaka M, Ohba Y, Miyazaki T, Imamura M, Takaoka A. 2011. ZAPS is a potent stimulator of signaling mediated by the RNA helicase RIG-I during antiviral responses. Nat Immunol 12:37–44. 10.1038/ni.1963.

9. Nguyen LP, Aldana KS, Yang E, Yao Z, Li MMH. 2023. Alphavirus Evasion of Zinc Finger Antiviral Protein (ZAP) Correlates with CpG Suppression in a Specific Viral nsP2 Gene Sequence. Viruses 15:830. doi: 10.3390/v15040830.

10. Law LMJ, Razooky BS, Li MMH, You S, Jurado A, Rice CM, MacDonald MR. 2019. ZAP’s stress granule localization is correlated with its antiviral activity and induced by virus replication. PLoS Pathog 15:e1007798. 10.1371/journal.ppat.1007798.

11. McPherson RL, Abraham R, Sreekumar E, Ong S, Cheng S, Baxter VK, Kistemaker HAV, Filippov DV, Griffin DE, Leung AKL. 2017. ADP-ribosylhydrolase activity of Chikungunya virus macrodomain is critical for virus replication and virulence. Proc Natl Acad Sci U S A 114:1666– 1671. 10.1073/pnas.1621485114.

12. Jayabalan AK, Adivarahan S, Koppula A, Abraham R, Batish M, Zenklusen D, Griffin DE, Leung AKL. 2021. Stress granule formation, disassembly, and composition are regulated by alphavirus ADP-ribosylhydrolase activity. Proc Natl Acad Sci U S A 118:e2021719118. doi: 10.1073/pnas.2021719118.

